# Coordinated Tuning of Ionizable Lipids and Formulation Redirects mRNA Vaccines Toward Lymphoid-Specific CD4^+^ T Cell Immunity

**DOI:** 10.64898/2026.04.16.719092

**Authors:** Yeji Lee, Yelim Choi, Seongryong Kim, Jinah Yeo, Junghwa Lee, Eun Hye Jeong, Jeong Hun Kwak, Min-Sung Kang, Ha-Eun Hong, Ok-Hee Kim, Yun-Ho Hwang, Jong-Eun Park, Eui Ho Kim, Say-June Kim, You-Jin Kim, Hyukjin Lee

## Abstract

The global success of mRNA vaccines has underscored the pivotal role of lipid nanoparticles (LNPs), yet how subtle chemical variations in ionizable lipids and their formulation parameters orchestrate complex immune landscapes remains largely elusive. Here, we report a novel ionizable lipid, N4Z, and demonstrate that its distinct chemical signature selectively intensifies early innate immune programs compared to its structural analogue, N4Y. Single-cell transcriptomic profiling at the injection site reveals that N4Z-based LNPs uniquely prime inflammatory and type I interferon-related transcriptional programs, accompanied by a rapid influx of B and CD4^+^ T cells. Beyond lipid chemistry, we show that formulation-level tuning, that is independent of the ionizable lipid structure, can reshape the systemic biodistribution from hepatic dominance toward lymphoid tissues. This optimization substantially enhances macrophage-associated antigen expression, which in turn amplifies polyfunctional CD4^+^ T cell responses, T follicular helper cell differentiation, and germinal center reactions. Our findings establish that the coordinated interplay between lipid engineering and formulation design provides a programmable platform for precision mRNA vaccination, achieving superior protective efficacy and neutralizing activity over clinically validated benchmarks.

## 1. Introduction

Messenger RNA (mRNA) vaccines have demonstrated remarkable clinical success, most notably during the COVID-19 pandemic, establishing mRNA as a versatile platform for vaccination and therapeutic development [1–4]. Central to this success is the lipid nanoparticle (LNP) delivery system, which protects mRNA from degradation and enables efficient intracellular delivery [5–7]. Accumulating evidence suggests that LNPs are not merely passive carriers but actively contribute to the magnitude and quality of immune responses elicited by mRNA vaccines [8, 9].

Among LNP components, ionizable lipids play a critical role by facilitating endosomal escape through pH-dependent protonation and largely determine transfection efficiency, biodistribution, and tolerability [10–13]. Beyond delivery efficiency, accumulating evidence indicates that ionizable lipid nanoparticles actively function as built-in adjuvants, inducing robust innate immune responses characterized by the production of inflammatory cytokines and type I interferons [14]. Recent studies have shown that LNP components can trigger IL-1– and IL-6–dependent inflammatory pathways, as well as type I interferon signaling at the injection site, which are critical for the induction of T follicular helper cells, germinal center reactions, and cellular immune responses [15–17].

In addition, formulation parameters such as lipid composition, particle size, and lipid ratios have been shown to critically influence cellular tropism, biodistribution, and the magnitude and quality of innate immune activation, thereby shaping downstream adaptive immune responses [18, 19]. Recent studies demonstrate that modulation of phospholipid species and PEG-lipid ratios can alter LNP biodistribution, immune cell uptake, and local inflammatory signaling, ultimately biasing vaccine-elicited immunity toward humoral or cellular responses [20]. Notably, systematic screening of LNP formulations has revealed that tuning helper lipid identity and component ratios can differentially program CD4⁺ T helper cell responses, including Th1- and Th2-skewed immunity, and enhance antigen presentation by antigen-presenting cells (APCs) [21].

Macrophages and other myeloid cells have emerged as key cellular targets of mRNA/LNP vaccines, serving not only as major sites of mRNA uptake and protein expression but also as central regulators of cytokine production and antigen presentation. Recent *in vivo* trafficking studies have shown that following intramuscular administration, mRNA/LNPs are preferentially taken up by myeloid populations at the injection site and in draining lymph nodes, where macrophages contribute to local innate immune activation and shape downstream adaptive immune responses [22]. In particular, lymph node–resident macrophages have been identified as critical drivers of type I interferon–dependent inflammatory programs that promote antigen-presenting cell maturation and T cell priming [23]. Consistent with this framework, growing evidence indicates that endogenous antigen expression within APCs plays a critical role in shaping CD4⁺ T cell responses. Antigen synthesized directly within APCs can be processed and presented via MHC class II pathways more efficiently than exogenously acquired antigen, resulting in enhanced CD4⁺ T cell priming and downstream humoral immunity [24].

Here, we address this gap by developing a novel ionizable lipid, N4Z, and systematically comparing its immunological properties with a structurally related analogue, N4Y, using a combination of *in vivo* immunological profiling and single-cell transcriptomics. We further investigate how formulation-level tuning of N4Z LNPs, independent of ionizable lipid structure, can modulate macrophage-associated antigen expression, reshape biodistribution, and amplify CD4⁺ T cell–dependent adaptive immunity. Together, this study establishes a framework for rational optimization of both ionizable lipid chemistry and formulation parameters to program innate and adaptive immune responses in mRNA vaccination.

## 2. Results

### 2.1. Rational Design and Physicochemical Profiling of an Ionizable Lipid Library

To engineer mRNA vaccine platforms with enhanced potency, we implemented a combinatorial synthesis of ionizable lipids and their formulations for systematic evaluation, focusing on the ionizable lipid as the primary determinant of both intracellular delivery kinetics and adjuvant-like immunological activity (**Figure 1**). Our synthetic strategy employed a modular approach where three distinct amine head groups (N1, N4, and N6) were combinatorially paired with three hydrophobic tails (X, Y, and Z) using a biodegradable 10-bromodecanoic acid linker. This design yielded a library of nine ionizable lipid candidates (**Figure 2a**), providing a structural framework to evaluate how subtle chemical variations orchestrate the interplay between delivery efficiency and immune programming. Structural confirmation of the representative lipid N4Z was performed by LC–MS (**Figure S1**), NMR spectroscopy (**Figure S2**), and HPLC analysis (**Figure S3**).

**Figure 1.**
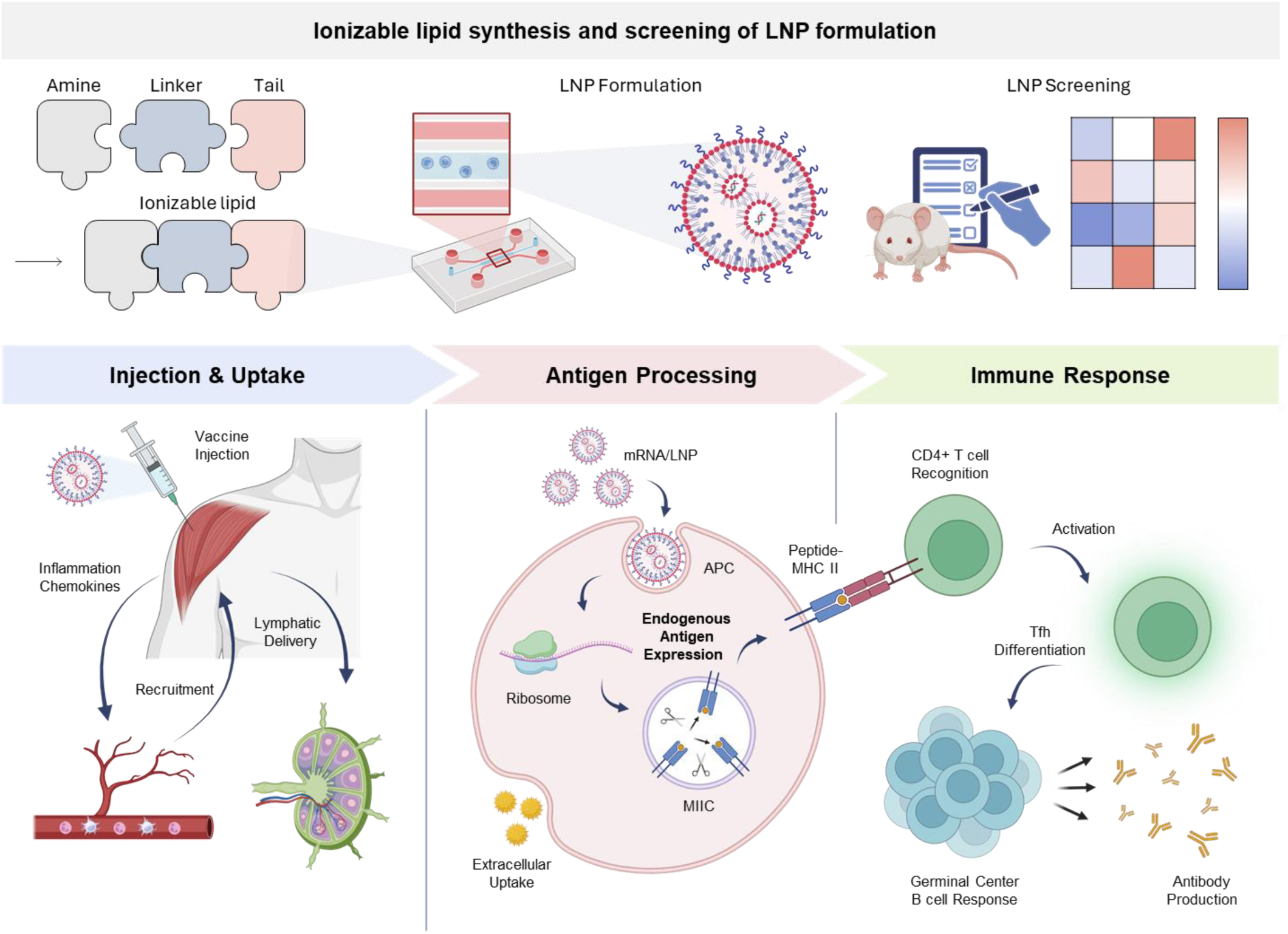
Integrated approach for the combinatorial synthesis of ionizable lipids and the systematic evaluation of LNP formulation. (Top) The LNP design and screening strategy illustrates the combinatorial synthesis of an ionizable lipid library by pairing various amine head groups and hydrophobic tails. These candidates are then formulated into LNPs using microfluidic mixing and subjected to systematic in vivo screening to identify lead ionizable lipids with optimal delivery efficiency. (Bottom) Following intramuscular administration, LNP–encapsulated mRNA is delivered to local tissues and draining lymph nodes. Internalized mRNA is translated into antigen, which is processed and presented by antigen-presenting cells (APCs) via MHC pathways. This process activates CD4^+^ T cells, promotes T follicular helper (Tfh) cell differentiation, and induces germinal center B cell responses, ultimately leading to robust antibody production.

**Figure 2.**
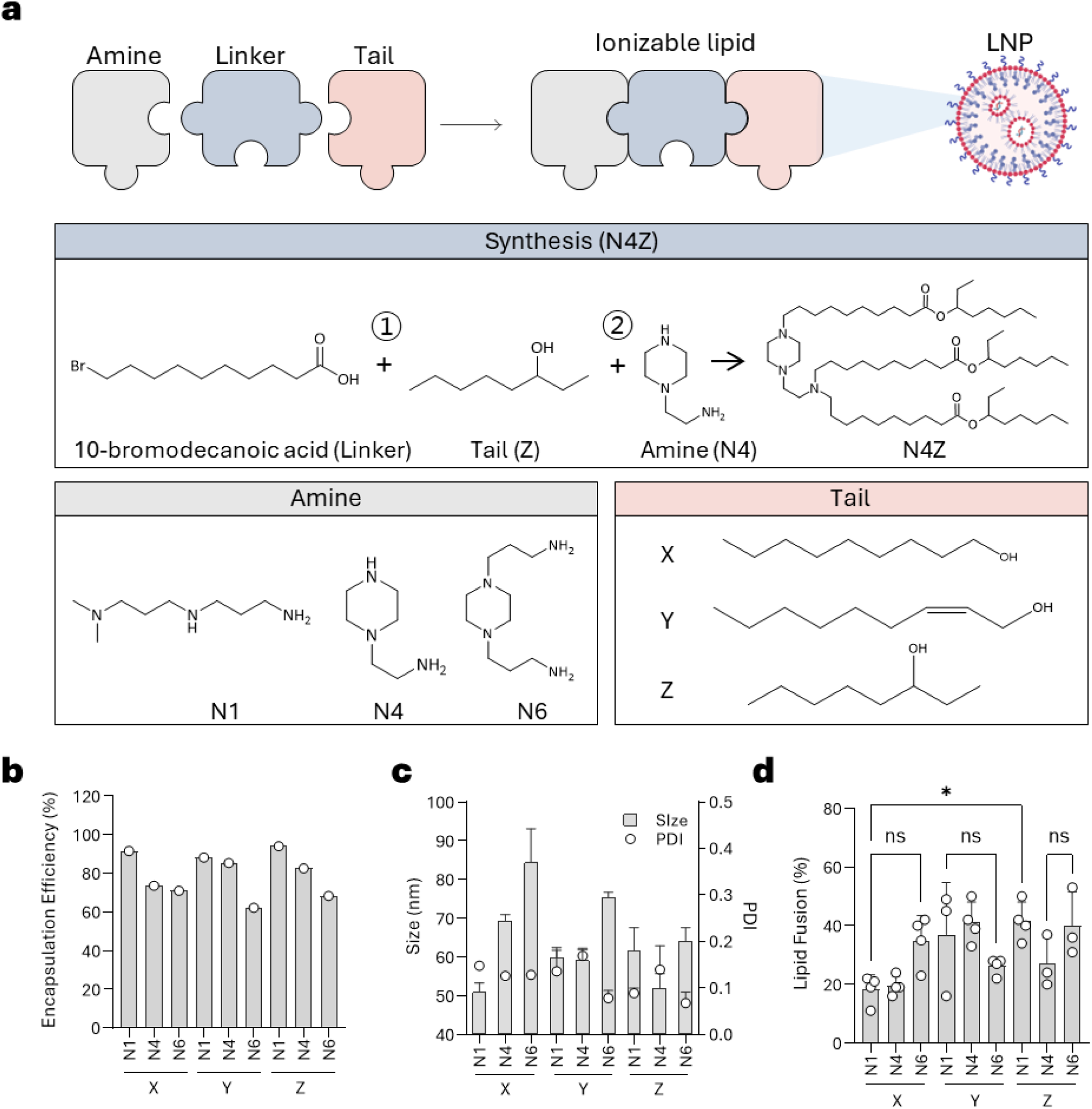
Design and physicochemical characterization of an ionizable lipid library. (a) Synthetic scheme of the ionizable lipid library comprising three amine head groups (N1, N4, N6) and three hydrophobic tails (X, Y, Z), generating nine candidates formulated into LNPs. (b) mRNA encapsulation efficiency. (c) Number-weighted hydrodynamic diameter and polydispersity index (PDI). (d) Lipid fusion efficiency assessed by an NBD-PE/Rhodamine-PE FRET-based lipid mixing assay using endosome-mimicking liposomes. Statistical significance was determined by one-way ANOVA followed by Tukey’s multiple comparisons test. Data are presented as mean ± SEM.

These ionizable lipids were formulated into LNPs using a microfluidic mixing platform, and their physicochemical properties were systematically evaluated. mRNA encapsulation efficiencies ranged from 62.2% to 94.1%, following a general trend of N1 > N4 > N6 (**Figure 2b**). Number-weighted hydrodynamic diameters ranged from approximately 51 to 84 nm with a size trend of N1 < N4 < N6, and all formulations maintained narrow size distributions (PDI < 0.2) (**Figure 2c**). Lipid fusion efficiencies, assessed using endosome-mimicking liposomes incorporating an NBD-PE/Rhodamine-PE FRET pair [25], ranged from approximately 18% to 42%, with Y- and Z-type tail-containing LNPs showing higher fusogenicity compared with X-type formulations (**Figure 2d**).

### 2.2. *In Vivo* Screening Identifies Lead Ionizable Lipids for Intramuscular mRNA Delivery

To identify ionizable lipid candidates with optimal intramuscular mRNA delivery efficiency, nine LNP formulations were evaluated *in vivo* using a luciferase reporter system (**Figure 3a**). C57BL/6N mice were intramuscularly injected with LNPs encapsulating firefly luciferase (fLuc) mRNA (0.05 mg/kg), and luciferase expression was assessed at 6 h post-injection using *in vivo* bioluminescence imaging.

**Figure 3.**
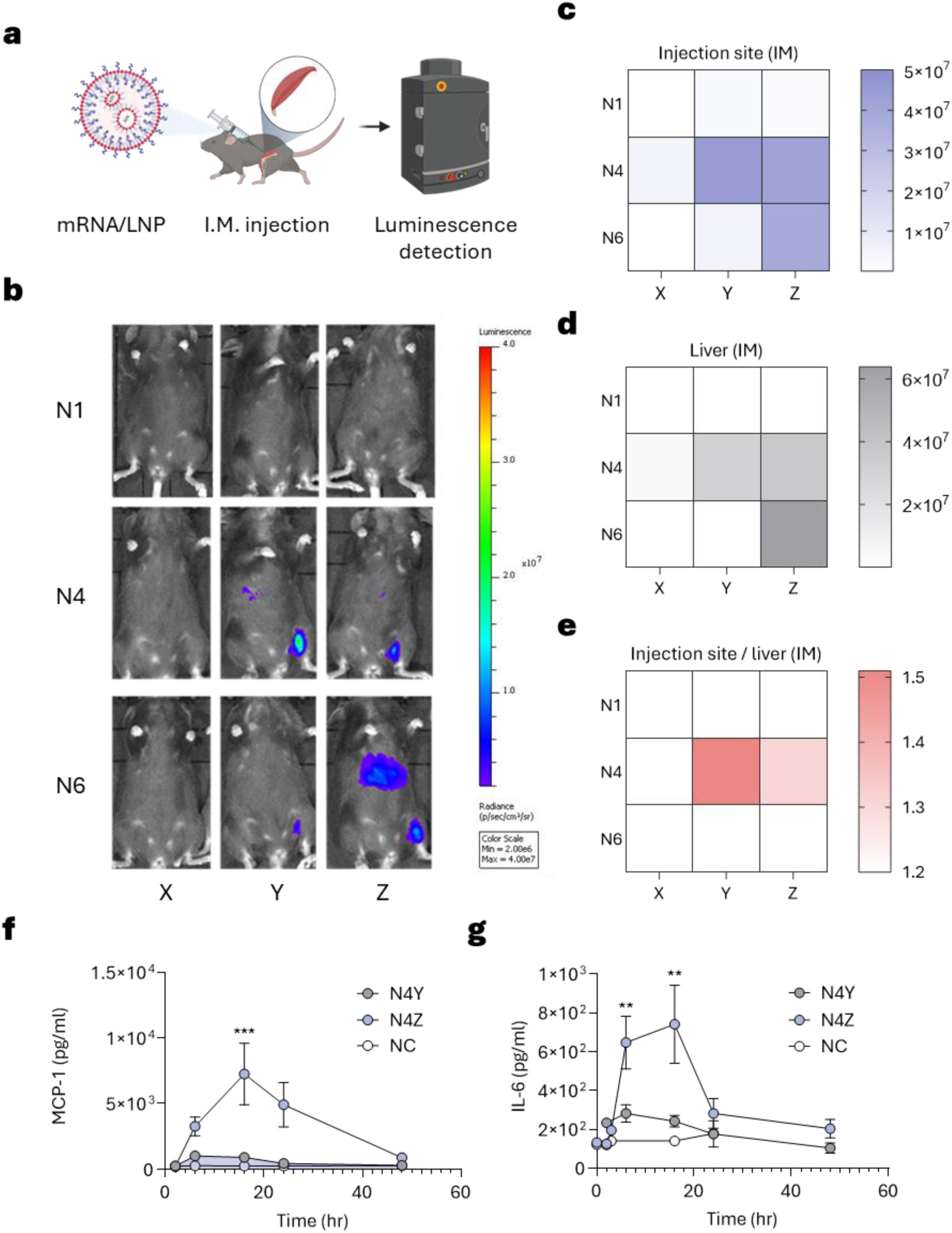
*In vivo* screening of ionizable lipid nanoparticles and early innate immune responses following intramuscular administration. (a) Experimental scheme for *in vivo* screening. C57BL/6N mice were intramuscularly administered fLuc mRNA-loaded LNPs (0.05 mg/kg) and bioluminescence was assessed at 6 h post-injection by IVIS imaging. (b) Representative *in vivo* IVIS bioluminescence images of nine LNP formulations. (c) Quantification of bioluminescence signals at the injection site. (d) Quantification of bioluminescence signals in the liver. (e) Injection site-to-liver bioluminescence ratio (n = 2 per group). N4Y and N4Z were selected as lead candidates based on high injection site expression and minimal hepatic off-target signal. (f, g) Time-dependent serum levels of MCP-1 (f) and IL-6 (g) following intramuscular administration of N4Y- or N4Z-based LNPs encapsulating SARS-CoV-2 spike mRNA (0.5 mg/kg) in BALB/c mice (n = 4 per group). Statistical significance was determined by unpaired two-tailed Student’s t-test at each time point. Data are presented as mean ± SEM. *p < 0.05, **p < 0.01, ***p < 0.001.

Bioluminescence signals at the injection site varied substantially across formulations (**Figure 3b, c**). N1-series and N6X-based LNPs showed comparatively low expression levels, whereas N4Y-, N4Z-, and N6Z-based LNPs exhibited the highest signals (approximately 4.4×10⁷, 4.2×10⁷, and 3.8×10⁷ p/s, respectively), suggesting that the combination of N4 head group with Y- or Z-type tails particularly favors intramuscular mRNA delivery. However, N6Z LNPs also showed substantial luciferase expression in the liver (approximately 6.4×10⁷ p/s), resulting in an injection site-to-liver ratio below 1 (approximately 0.31) (**Figure 3d, e**). Hepatic off-target distribution is undesirable in the vaccine context, as it may reduce local immune activation at the injection site while increasing the risk of systemic adverse effects. In contrast, N4Y and N4Z maintained injection site-dominant expression profiles with site-to-liver ratios of approximately 1.51 and 1.30, respectively, consistent with the localized intramuscular expression pattern considered favorable for driving efficient innate immune activation and downstream adaptive responses. Based on these results, N6Z was excluded from further evaluation, and N4Y and N4Z were selected as lead ionizable lipid candidates for subsequent studies.

### 2.3. N4Z LNPs Induce Enhanced Early Cytokine and Chemokine Responses *In Vivo*

Innate immune activation, including early cytokine and chemokine induction, plays a critical role in shaping vaccine immunogenicity and promoting adaptive immune responses [26, 27]. To evaluate whether lead LNP formulations function as immunostimulatory vaccine delivery systems, *in vivo* cytokine and chemokine responses were assessed following mRNA/LNP administration.

BALB/c mice were intramuscularly injected with LNPs encapsulating SARS-CoV-2 spike mRNA (0.5 mg/kg), and serum samples were collected at multiple time points post-injection. Levels of the chemokine MCP-1 (**Figure 3f**) and cytokine IL-6 (**Figure 3g**) were quantified by ELISA. Both formulations induced measurable increases in MCP-1 and IL-6; however, N4Z LNPs elicited significantly higher levels compared with N4Y. MCP-1 levels peaked at 16 h post-injection in the N4Z group (approximately 7,260 pg/mL) compared with N4Y (approximately 900 pg/mL, p < 0.001), and IL-6 levels were also significantly elevated at 6 h and 16 h post-injection (p < 0.01).

Both cytokine and chemokine levels declined gradually toward baseline by 48 h, indicating a transient inflammatory profile. These kinetics suggest that N4Z promotes robust early innate immune activation without prolonged systemic inflammation, a feature considered favorable for effective vaccine adjuvanticity.

### 2.4. Single-Cell Transcriptomic Analysis Reveals Enhanced Inflammatory and Interferon Programs Induced by N4Z LNPs at the Injection Site

To investigate early cellular responses at the injection site, BALB/c mice were intramuscularly administered N4Y- or N4Z-based LNPs encapsulating SARS-CoV-2 spike mRNA (0.25 mg/kg), and muscle tissues at the injection site were collected 16 h post-injection for single-cell RNA sequencing (scRNA-seq) analysis (**Figure 4a**). Analysis of cell-type–resolved transcript detection revealed that both formulations enabled measurable spike mRNA detection across multiple stromal and immune cell populations, including fibroblasts, macrophages, monocytes, and endothelial cells (**Figure 4b, c**). Notably, spike mRNA was robustly detected in fibroblasts for both LNP formulations, indicating that stromal cells represent a major site of local mRNA presence following intramuscular administration.

**Figure 4.**
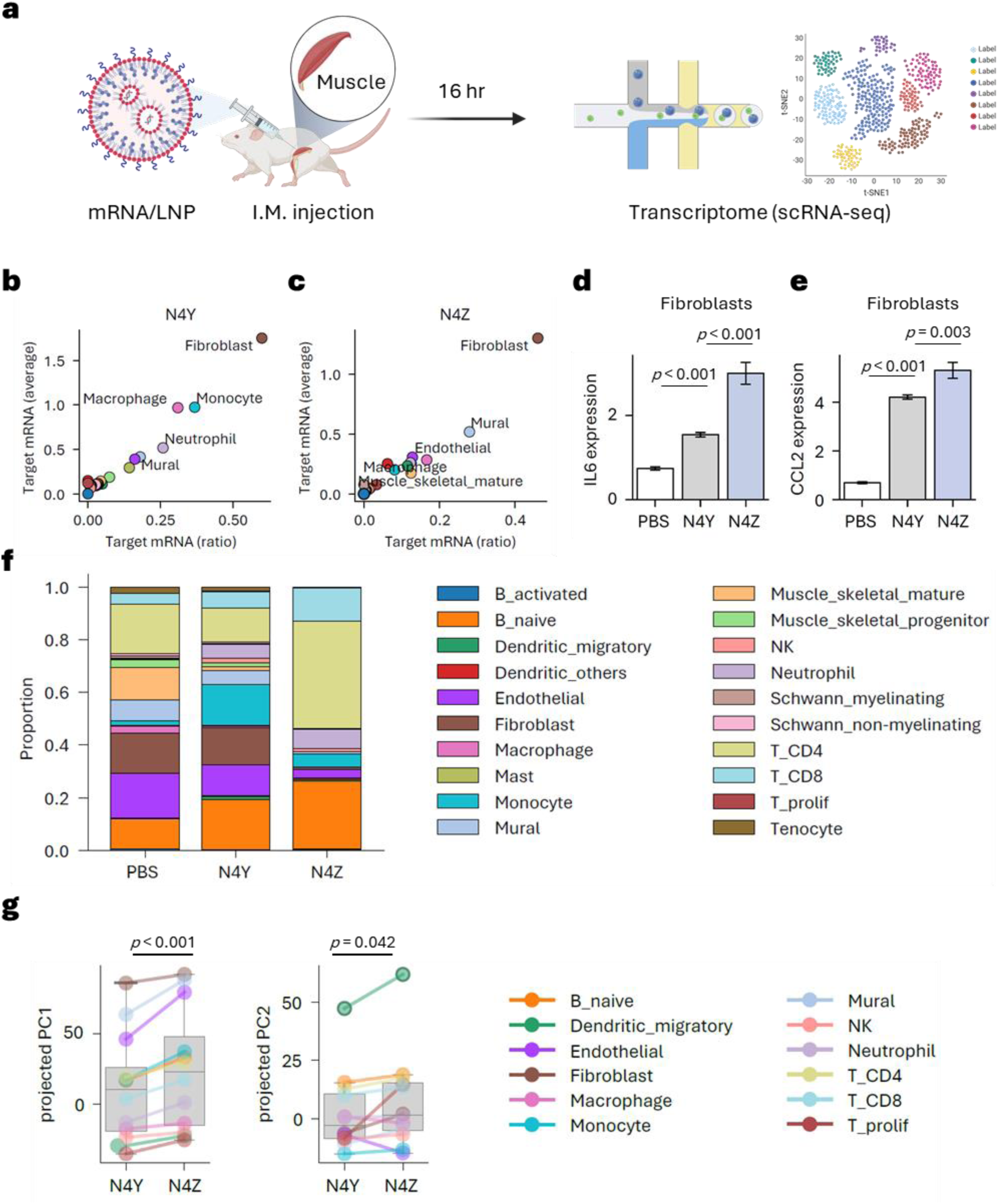
Single-cell transcriptomic profiling at the injection site following mRNA/LNP administration. (a) Experimental scheme. BALB/c mice were intramuscularly administered N4Y- or N4Z-based LNPs encapsulating SARS-CoV-2 spike mRNA (0.25 mg/kg), and muscle tissues at the injection site were collected 16 h post-injection for scRNA-seq analysis. (b, c) Scatter plots showing the proportion and average expression of spike mRNA detected across cell types following administration of N4Y- (b) and N4Z-based (c) LNPs. (d, e) Relative expression of *Il6* (d) and *Ccl2* (e) transcripts in injection site fibroblasts across PBS, N4Y, and N4Z conditions. Statistical significance was determined by one-way ANOVA followed by Tukey’s multiple comparisons test. (f) Cellular composition of profiled cells at the injection site across conditions. (g) Projected PC1 and PC2 scores of aggregated cell-type transcriptional profiles comparing N4Y and N4Z conditions, where PC1 reflects inflammatory activation and PC2 captures an mRNA/LNP-associated transcriptional program enriched for type I interferon–related gene signatures. Statistical significance was determined by Wilcoxon signed-rank test. Data are presented as mean ± SEM. \**p* < 0.05, \*\**p* < 0.01, \*\*\**p* < 0.001.

To assess local inflammatory signaling, expression of cytokine and chemokine transcripts was examined at single-cell resolution. Fibroblasts at the injection site showed significantly elevated *Il6* and *Ccl2* transcript levels following N4Z administration compared with N4Y (**Figure 4d, e**), consistent with enhanced innate immune activation observed *in vivo*. Analysis of cellular composition at the injection site revealed notable differences between conditions (**Figure 4f**). While major stromal populations such as muscle cells remained comparable across groups, N4Z administration was associated with a higher proportion of B naïve cells and CD4⁺ T cells compared with N4Y and PBS controls, alongside reductions in endothelial and monocyte fractions. These compositional shifts, particularly the increase in B naïve cells and CD4⁺ T cells in the N4Z group, suggest that heightened inflammatory signaling may influence local immune cell dynamics as early as 16 h post-injection.

To further characterize global transcriptional responses, principal component analysis was performed on aggregated cell-type transcriptional profiles (**Figure 4g**). Gene set enrichment analysis revealed that PC1 was dominated by cytokine- and inflammation-related pathways, while PC2 was strongly enriched for type I interferon signaling and antiviral response programs (**Figure S4**), indicating that PC1 reflects inflammatory activation and PC2 captures an mRNA/LNP-driven transcriptional program enriched for type I interferon–related gene signatures. N4Z-treated samples showed consistently higher projected PC1 scores across most cell types and significantly elevated projected PC2 scores compared with N4Y (PC1, p < 0.001; PC2, p = 0.042), with particularly pronounced increases observed in dendritic cell populations.

### 2.5. N4Z-based mRNA vaccination enhances adaptive immune responses and confers protective efficacy

To evaluate adaptive immune responses, BALB/c mice were immunized by intramuscular injection with SARS-CoV-2 spike mRNA at a dose of 0.25 mg/kg encapsulated in N4Y or N4Z LNPs using a prime–boost regimen with a three-week interval (**Figure 5a**). Spike-specific IgG levels, measured three weeks after booster immunization, were markedly elevated in the N4Z group compared with N4Y (approximately 6,642,040 vs. 124,540, respectively (**Figure 5b**). Neutralizing antibody responses assessed by PRNT₅₀ were significantly higher in N4Z-immunized mice compared with the N4Y group (approximately 4,870 vs. 2,605, respectively), indicating substantially enhanced humoral immunity (**Figure 5c**). Antigen-specific cellular immunity assessed by ELISpot showed that N4Z vaccination induced modestly stronger T cell responses compared with N4Y (approximately 138 vs. 107 spots, respectively) (**Figure 5d**).

**Figure 5.**
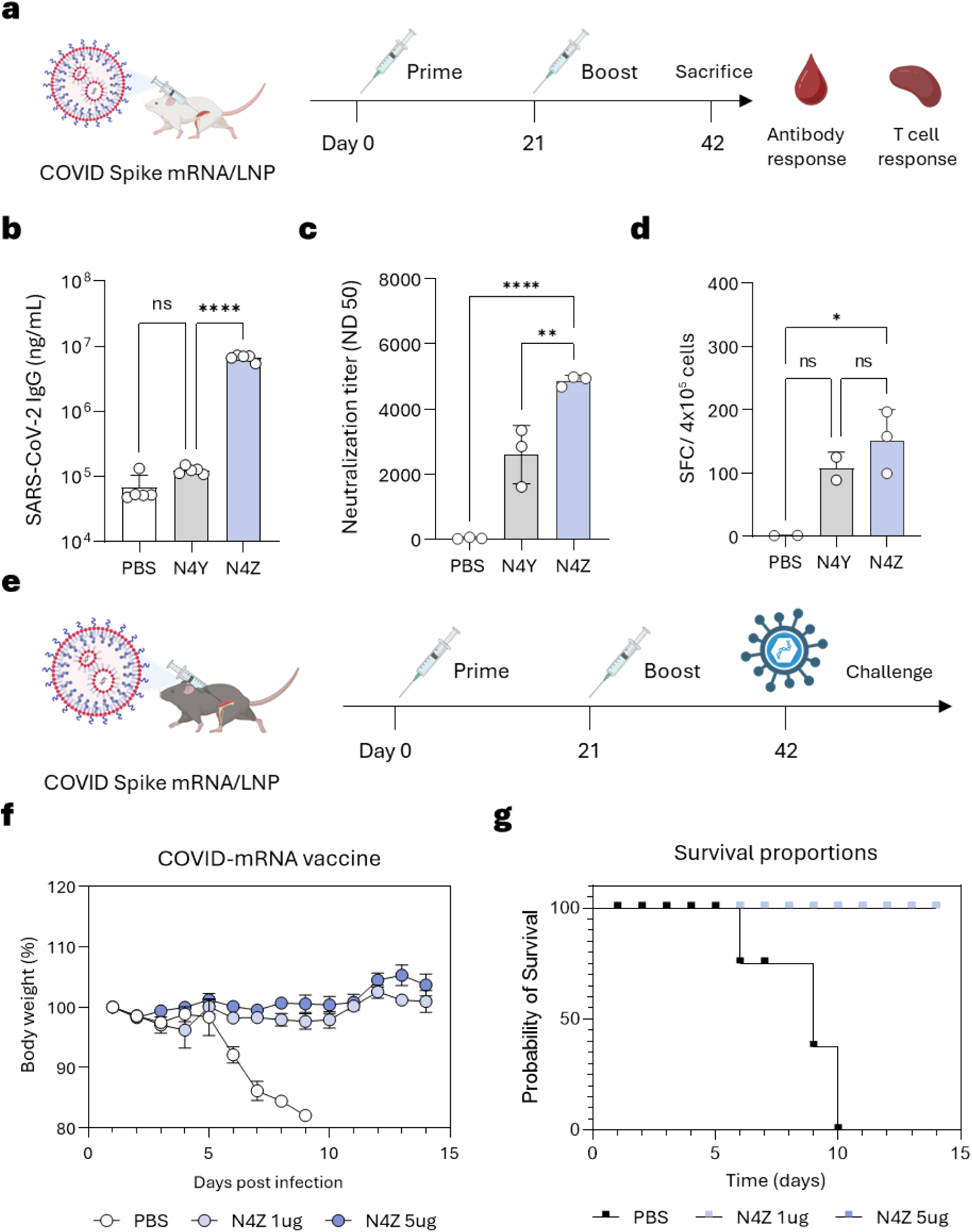
Adaptive immune responses and protective efficacy of N4Y- and N4Z-based mRNA vaccines. (a) Vaccination scheme. BALB/c mice were immunized intramuscularly with SARS-CoV-2 spike mRNA-loaded LNPs (0.25 mg/kg) following a prime–boost regimen (day 0 and day 21). Blood samples and splenocytes were collected three weeks after the booster dose for antibody and T cell analysis. (b) Antigen-specific IgG levels at three weeks post-boost (n = 5 per group). (c) Neutralizing antibody titers assessed by PRNT₅₀ at three weeks post-boost (n = 3 per group). (d) Antigen-specific T cell responses assessed by IFN-γ ELISpot (n= 2–3 per group). (e) Viral challenge scheme. K18-hACE2 transgenic mice were immunized following the same prime–boost regimen and challenged with SARS-CoV-2 three weeks after the booster dose. (f) Body weight changes following viral challenge (n = 5 per group). (g) Survival curves following viral challenge (n = 5 per group). Statistical significance for (b–d) was determined by one-way ANOVA followed by Tukey’s multiple comparisons test. Survival curves were compared using the log-rank (Mantel-Cox) test. Data are presented as mean ± SEM. *p < 0.05, **p < 0.01, ***p < 0.001.

To determine whether these immune responses translated into protection, K18-hACE2 transgenic mice were immunized following the same prime–boost regimen and subsequently challenged with SARS-CoV-2 three weeks after the booster dose (**Figure 5e**). N4Z-vaccinated animals maintained stable body weight throughout the observation period, remaining within approximately 97–105% of their initial body weight at both 1 μg and 5 μg doses, whereas PBS control animals exhibited progressive weight loss following infection (**Figure 5f**). Importantly, all N4Z-vaccinated mice survived the viral challenge regardless of dose, achieving 100% survival, while PBS control animals showed progressive mortality beginning at day 6 post-challenge (**Figure 5g**). Booster dosing markedly increased neutralizing antibody titers compared with the priming response, confirming robust recall immunity induced by N4Z vaccination (**Figure S5**). Collectively, these results demonstrate that N4Z-based mRNA vaccination induces potent humoral and cellular immunity and provides effective protection against viral challenge.

### 2.6. Safety evaluation of N4Z lipid nanoparticles

To assess the *in vivo* safety profile of N4Z LNPs, BALB/c mice were intramuscularly administered N4Z LNPs at doses of 0.2 or 1.0 mg/kg, and serum biochemical markers, histological analysis, and body weight changes were evaluated at 24 h post-injection. Liver function markers (AST: ∼170–210 U/L; ALT: ∼28 U/L) and kidney function markers (BUN: ∼17–20 mg/dL; creatinine: ∼0.15 mg/dL) remained within ranges comparable to untreated controls across both doses, indicating no overt hepatic or renal toxicity (**Figure S6a**). Histological examination of liver tissue further confirmed the absence of noticeable pathological changes or inflammatory lesions in N4Z-treated groups (**Figure S6b**). Consistent with these findings, body weight remained stable across all groups (approximately 20–22 g), with no significant changes observed relative to pre-dose measurements (**Figure S6c**). Collectively, these results demonstrate that N4Z LNPs exhibit a favorable safety profile without detectable acute toxicity under the tested conditions.

### 2.7. Formulation-level optimization enhances macrophage-associated mRNA expression and reshapes biodistribution

Macrophages are key cellular targets of mRNA/LNP vaccines and play critical roles in antigen expression and downstream immune programming [22, 23]. Consistent with this, endogenous antigen expression within antigen-presenting cells has been shown to promote efficient MHC class II presentation and enhance CD4⁺ T cell priming and humoral immunity [24]. Given the emerging role of macrophages and antigen-presenting cells as central drivers of CD4⁺ T cell–dependent immunity, we hypothesized that further tuning of LNP formulation parameters could enhance APC-associated antigen expression and thereby improve adaptive immune responses.

To further improve the performance of N4Z-based LNPs, formulation parameters were optimized while maintaining identical ionizable lipid structure (**Figure 6a**). Specifically, the helper lipid composition was adjusted by increasing DOPE and reducing cholesterol to modulate membrane fusion and potentially limit ApoE-mediated hepatic uptake, and the PEG-lipid component was switched from C16-PEG ceramide to DMG-PEG [28]. DMG-PEG exhibits faster PEG shedding kinetics compared with C16-PEG ceramide, facilitating more rapid exposure of the nanoparticle surface after administration [29]. The overall PEG-lipid content was also reduced from 1.5% to 0.5% to further promote cellular interactions by minimizing surface shielding [30, 31]. Additionally, the lipid-to-RNA weight ratio was increased from 10 to 15. These combined changes resulted in a marked increase in particle size from approximately 53 nm (N4Z) to approximately 120 nm (N4Z-opt) (**Figure 6c**), likely reflecting decreased lipid packing density. Given that macrophage uptake of nanoparticles has been reported to be size-dependent, this increase in particle size may have further contributed to enhanced macrophage-associated mRNA expression [32]. Notably, N4Z-opt maintained a high RNA encapsulation efficiency of approximately 90.6% (**Figure 6b**), comparable to the original N4Z formulation (approximately 95.4%), and exhibited a lower polydispersity index (PDI ∼0.14 vs. ∼0.18) (**Figure 6d**), indicating a uniform and well-defined nanoparticle population.

**Figure 6.**
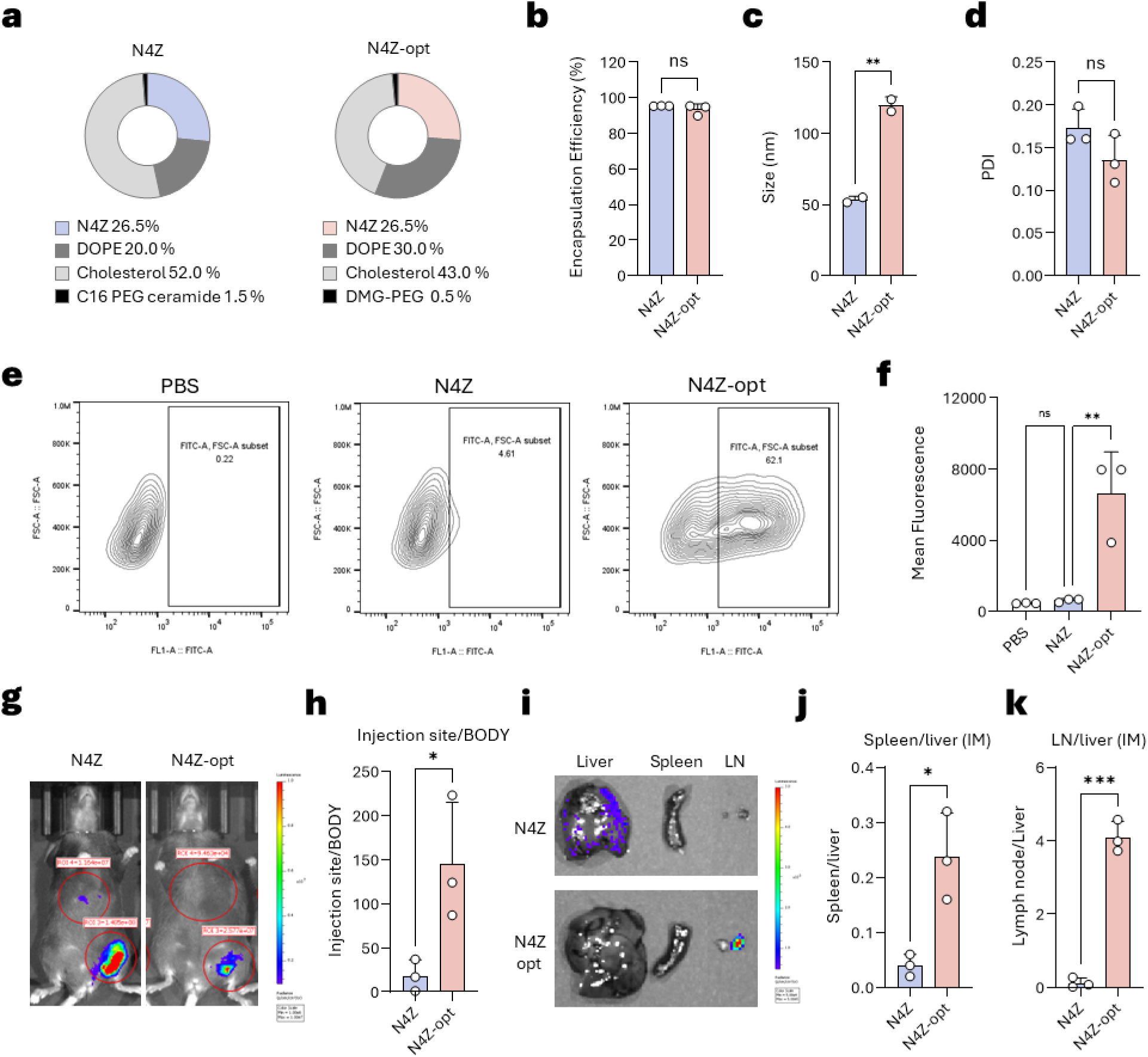
Formulation-level optimization of N4Z lipid nanoparticles and biodistribution. (a) Schematic comparison of lipid compositions of N4Z and optimized N4Z (N4Z-opt) formulations. (b) mRNA encapsulation efficiency (n = 3). (c) Hydrodynamic diameter (n = 3). (d) Polydispersity index (PDI) (n = 3). (e) Representative flow cytometry contour plots showing GFP expression in RAW 264.7 macrophages following delivery of PBS, N4Z, and N4Z-opt. (f) Mean fluorescence intensity (MFI) of GFP expression in RAW 264.7 macrophages (n = 3). (g) Representative *in vivo* IVIS bioluminescence images following intramuscular administration of fLuc mRNA-loaded N4Z or N4Z-opt LNPs. (h) Injection site-to-body bioluminescence ratio (n = 3). (i) Representative *ex vivo* organ images showing bioluminescence in liver, spleen, and lymph node. (j) Spleen-to-liver bioluminescence ratio (n = 3). (k) Lymph node-to-liver bioluminescence ratio (n = 3). Statistical significance for (b–d) was determined by unpaired two-tailed Student’s *t*-test. Statistical significance for (f, h, j, k) was determined by one-way ANOVA followed by Tukey’s multiple comparisons test. Data are presented as mean ± SEM. \**p* < 0.05, \*\**p* < 0.01, \*\*\**p* < 0.001.

We next evaluated cellular transfection efficiency using GFP mRNA in RAW 264.7 macrophage cells. Flow cytometry analysis revealed a markedly increased mean fluorescence intensity (MFI) following delivery of N4Z-opt compared with the original N4Z formulation (approximately 7,929 vs. 614, respectively), representing an approximately 13-fold enhancement in macrophage transfection efficiency (**Figure 6e, f**). These findings demonstrate that formulation-level optimization substantially enhances macrophage-associated mRNA delivery without modifying the ionizable lipid structure.

### 2.8. Formulation optimization enhances local expression and redirects biodistribution toward lymphoid tissues

To determine whether formulation optimization influences *in vivo* expression and biodistribution, bioluminescence imaging was performed following intramuscular administration of fLuc mRNA–loaded LNPs in C57BL/6N mice. Representative IVIS images demonstrated visibly stronger luminescence at the injection site in N4Z-opt–administered mice compared with N4Z (**Figure 6g**). Quantification confirmed that N4Z-opt exhibited a significantly higher injection site–to–body bioluminescence ratio compared with N4Z (approximately 145 vs. 18, p < 0.05), indicating markedly improved localization of mRNA expression at the site of administration (**Figure 6h**). This enhanced local expression is consistent with the improved macrophage transfection efficiency observed *in vitro*, suggesting that the formulation-driven increase in cellular uptake translates into more efficient mRNA expression at the injection site *in vivo*.

*Ex vivo* organ imaging further revealed a striking shift in biodistribution between the two formulations (**Figure 6i**). While N4Z showed predominant expression in the liver, consistent with ApoE-mediated hepatic uptake commonly observed with conventional LNP formulations, N4Z-opt exhibited markedly higher bioluminescence signals in the spleen and lymph nodes relative to the liver. Quantification confirmed significantly elevated spleen-to-liver (approximately 0.24 vs. 0.04, p < 0.05) and lymph node-to-liver ratios (approximately 3.97 vs. 0.11, p < 0.001) in the N4Z-opt group (**Figure 6j, k**). This redistribution toward lymphoid tissues likely reflects the combined effects of faster PEG shedding kinetics, reduced PEG density, and increased particle size in N4Z-opt, which together reduce ApoE-mediated hepatic targeting and promote preferential uptake by macrophages and antigen-presenting cells in immune-relevant compartments. Comprehensive *ex vivo* imaging demonstrated minimal bioluminescence signal in other major organs for both formulations (**Figure S7**), supporting the specificity of this biodistribution shift toward lymphoid tissues. Collectively, these results demonstrate that formulation-level optimization not only enhances local mRNA expression at the injection site but also redirects biodistribution toward immune-relevant lymphoid tissues, providing a mechanistic rationale for the enhanced adaptive immune responses demonstrated with N4Z-opt.

### 2.9. Optimized N4Z formulation enhances CD4⁺ T cell responses and humoral immunity

To evaluate whether formulation-level optimization improves adaptive immune responses, BALB/c mice were immunized intramuscularly with spike mRNA–loaded LNPs (0.25 mg/kg) following a prime–boost regimen (day 0 and day 21) (**Figure 7a**). Antigen-specific CD4⁺ T cell functionality was assessed by intracellular cytokine staining. N4Z-opt significantly increased the frequencies of IFN-γ⁺ (approximately 0.048% vs. 0.032%), TNF-α⁺ (approximately 0.145% vs. 0.084%), and IL-2⁺ (approximately 0.155% vs. 0.099%) CD4⁺ T cells compared with the original N4Z formulation (**Figure 7b–d**). In addition, N4Z-opt increased the proportion of polyfunctional CD4⁺ T cells capable of producing multiple cytokines simultaneously (approximately 0.109% vs. 0.069%) (**Figure 7e**), suggesting improved functional quality of vaccine-induced T cell responses. Representative flow cytometry plots of cytokine-producing CD4⁺ T cells from the spleen are shown in **Figure S8**.

**Figure 7.**
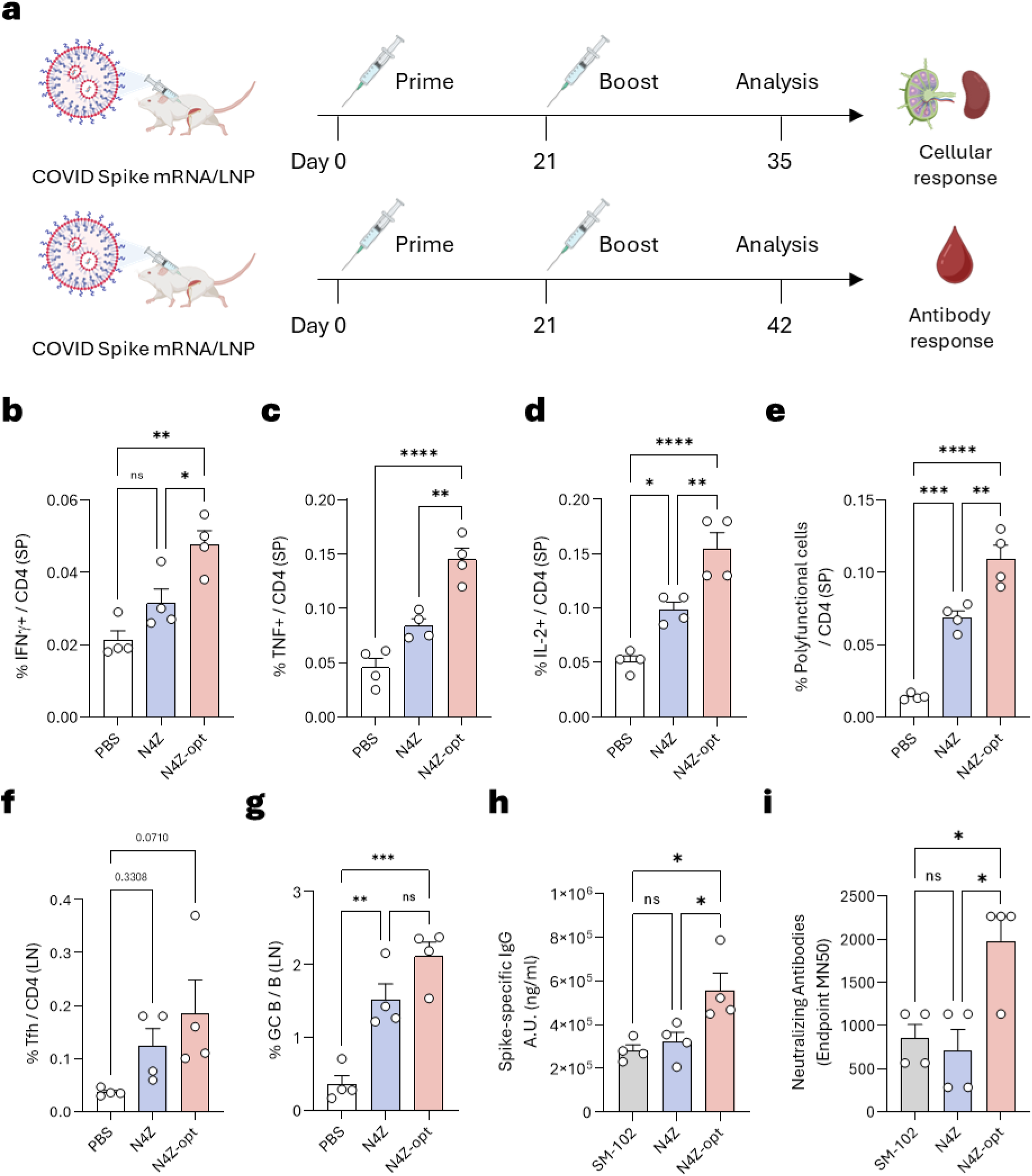
Enhanced CD4⁺ T cell and humoral immune responses induced by the optimized N4Z formulation. (a) Vaccination and analysis scheme. BALB/c mice were immunized intramuscularly with SARS-CoV-2 spike mRNA-loaded LNPs (0.25 mg/kg) following a prime–boost regimen (day 0 and day 21). Cellular immune responses were analyzed at day 35 (two weeks post-boost) following sacrifice, and antibody responses were assessed at day 42 (three weeks post-boost). (b–d) Frequencies of IFN-γ⁺ (b), TNF-α⁺ (c), and IL-2⁺ (d) cells among splenic CD4⁺ T cells (n = 4 per group). (e) Frequency of polyfunctional CD4⁺ T cells (n = 4 per group). (f) T follicular helper (Tfh) cell frequency in draining lymph nodes (n = 4 per group). (g) Germinal center (GC) B cell frequency in draining lymph nodes (n = 4 per group). (h) Antigen-specific IgG levels at day 42 (n = 4 per group). (i) Neutralizing antibody titers assessed by endpoint MN50 at day 42 (n = 4 per group). Statistical significance was determined by one-way ANOVA followed by Tukey’s multiple comparisons test. Data are presented as mean ± SEM. *p < 0.05, **p < 0.01, ***p < 0.001.

In contrast to CD4⁺ T cell responses, formulation-level optimization did not significantly alter antigen-specific CD8⁺ T cell frequencies or polyfunctional responses (**Figure S9**), indicating that N4Z-opt selectively enhances helper T cell–associated immunity without broadly amplifying cytotoxic T cell responses.

Given the critical role of CD4⁺ T cells in supporting germinal center formation, we next examined T follicular helper (Tfh) cells and germinal center (GC) B cells in draining lymph nodes. N4Z-opt significantly increased Tfh cell frequency (approximately 0.185% vs. 0.124%) and GC B cell responses (approximately 2.12% vs. 1.52%) compared with the original N4Z formulation (**Figure 7f, g**), suggesting improved support for B cell maturation and antibody development. Representative flow cytometry plots from draining lymph nodes are shown in **Figure S10**.

Consistent with these findings, antigen-specific humoral responses were substantially enhanced following vaccination with N4Z-opt. Spike-specific IgG levels at day 42 were significantly higher in the N4Z-opt group compared with both the original N4Z formulation and SM-102 (approximately 557,816 vs. 322,627 and 282,249, respectively) (**Figure 7h**). Neutralizing antibody titers assessed by MN50 at day 42 were also markedly elevated in the N4Z-opt group compared with N4Z and SM-102 (approximately 1,980 vs. 707 and 848, respectively) (**Figure 7i**), demonstrating that formulation-level optimization substantially enhanced both antibody magnitude and neutralizing capacity.

Collectively, these results demonstrate that formulation-level optimization of N4Z selectively amplifies CD4⁺ T cell functionality and germinal center responses, promotes stronger and more durable humoral immunity, and induces clinically relevant neutralizing activity, establishing N4Z-opt as an improved mRNA vaccine platform.

## 3. Conclusion

In this study, we developed a novel ionizable lipid, N4Z, and demonstrated that subtle structural variation in ionizable lipids can significantly influence both innate and adaptive immune responses in mRNA vaccination. Through systematic comparison with a closely related analogue, N4Y, we show that N4Z enhances early inflammatory and type I interferon–associated transcriptional programs, promotes stronger CD4⁺ T cell–dependent adaptive immunity, and improves protective efficacy without detectable acute toxicity. Furthermore, formulation-level optimization of N4Z reshaped biodistribution, enhanced macrophage-associated mRNA expression, and further amplified helper T cell and germinal center responses.

Ionizable lipids are primarily known for their role in facilitating endosomal escape and determining delivery efficiency; however, accumulating evidence suggests that they also function as key immunomodulatory components of mRNA/LNP vaccines. Our findings support this concept by demonstrating that a subtle modification in ionizable lipid structure can substantially alter early innate immune activation, accompanied by early shifts in local immune cell composition. Single-cell transcriptomic analysis revealed that N4Z selectively amplified inflammatory and type I interferon–related transcriptional programs at the injection site. These pathways are known to play critical roles in antigen-presenting cell activation, T follicular helper cell differentiation, and germinal center formation, which collectively contribute to the development of robust humoral immunity. Consistent with this framework, N4Z induced higher antigen-specific IgG levels and stronger neutralizing antibody responses compared with N4Y.

Another important observation from this study is that formulation-level tuning further enhanced the immunological performance of N4Z without altering its chemical structure. Specifically, modulation of formulation parameters increased macrophage-associated mRNA expression and shifted biodistribution toward immune-relevant tissues such as the spleen and lymph nodes. Macrophages and other myeloid populations have emerged as key regulators of mRNA vaccine–induced immune responses, serving as major sites of antigen expression and cytokine production. Enhanced antigen synthesis within antigen-presenting cells is known to improve MHC class II presentation and CD4⁺ T cell priming. Consistent with this mechanism, the optimized N4Z formulation selectively amplified CD4⁺ T cell functionality, increased T follicular helper and germinal center B cell responses, and promoted stronger and more durable humoral immunity, while having minimal impact on CD8⁺ T cell responses. Notably, N4Z-opt also demonstrated superior antibody magnitude and neutralizing activity compared with SM-102, a clinically validated ionizable lipid, highlighting the translational potential of this optimized platform. These findings highlight the importance of delivery-driven antigen expression in shaping helper T cell–dependent immune programming.

Safety is a critical consideration for next-generation ionizable lipids. Importantly, N4Z LNPs showed no detectable acute hepatic or renal toxicity, no significant histopathological abnormalities, and good systemic tolerability under the tested conditions. The transient cytokine profile observed following N4Z administration suggests effective innate immune activation without prolonged systemic inflammation, a balance that is considered favorable for vaccine adjuvanticity. Nevertheless, comprehensive long-term and dose-escalation safety studies will be required to fully evaluate the translational potential of this lipid platform.

While these findings highlight the robust potential of our platform in enhancing immune responses and protective efficacy, certain considerations are warranted to facilitate its clinical translation. Although we have demonstrated clear benefits in murine models, further studies are needed to account for species-specific differences in LNP biodistribution and immune activation that may occur in humans. Additionally, while our data suggest that macrophage-associated antigen expression plays a key role in improving CD4^+^ T cell immunity, direct causal validation through cell-specific targeting or depletion will further solidify this mechanism. Finally, expanding our structural-immunological mapping across a broader ionizable lipid chemical space will continue to refine and optimize our rational design principles.

In conclusion, our findings demonstrate that coordinated tuning of ionizable lipid chemistry and formulation parameters provides a powerful strategy to modulate innate immune activation, antigen-presenting cell–associated antigen expression, and CD4⁺ T cell–dependent adaptive immunity in mRNA vaccination. These results highlight the dual role of ionizable lipids as both delivery systems and immune modulators and provide design insights for next-generation mRNA vaccine platforms with improved immunogenicity and precision immune programming.

## 4. Experimental Section/Methods

### Materials

Amines, including 1-nonanol (8.06866), 3-octanol (W358118), cis-2-nonene-1-ol (W372005), N,N-dimethyldipropylenetriamine (550019), 1-(2-aminoethyl)piperazine (A55209), and 1,4-bis(3-aminopropyl)piperazine (239488), were purchased from TCI and Sigma-Aldrich. Lipid components, including 1,2-dioleoyl-sn-glycero-3-phosphoethanolamine (DOPE), 1,2-dimyristoyl-rac-glycero-3-methoxypolyethylene glycol-2000 (DMG-PEG, 880151P), and C16-PEG 2000 ceramide (880180), were obtained from Avanti Polar Lipids. Cholesterol (C8667) was purchased from Sigma-Aldrich (St. Louis, MO, USA).

D-luciferin (P1043) was purchased from Promega. RNA encapsulation efficiency was determined using the Quant-iT™ RiboGreen® assay (Life Technologies, USA). Reporter mRNAs, including EGFP (L-7201) and fLuc (L-7202), were obtained from TriLink BioTechnologies. SARS-CoV-2 spike protein mRNA (N1-methyl-pseudouridine modified; CS-0650147-1mg) was purchased from Chemscene. Mouse MCP-1 (BMS6005) and IL-6 (KMC0061) ELISA kits were obtained from Thermo Fisher Scientific (Waltham, MA, USA).

### Synthesis of biodegradable ionizable lipid library

Brominated acid linkers were added to 10 mL of dichloromethane (DCM) containing a mixture of 1-nonanol (X), 3-octanol (Y), cis-2-nonene-1-ol (Z), DIC, and DMAP. The reaction mixture was stirred at room temperature for 16 h. The resulting brominated ester intermediates were isolated using a solvent gradient of 0–10% ethyl acetate in hexane, followed by purification via column chromatography on a CombiFlash Rf system.

The purified carbon-tail intermediates were subsequently dissolved in 10 mL of acetonitrile (ACN) and reacted with the corresponding amine in the presence of DIPEA. The reaction was stirred at 80 °C for 72 h, after which the solvent was removed by evaporation at ambient temperature. The final product was purified by column chromatography on a CombiFlash Rf system using a gradient of 0–10% methanol in DCM.

### Preparation of LNPs

LNPs were prepared by mixing an ethanol phase containing lipid components with an aqueous RNA phase using a microfluidic mixing system (Ignite, PNI, Canada), following previously reported protocols [24]. Ionizable lipids, cholesterol, helper lipid (DOPE), and PEG lipids (C16-PEG 2000 ceramide and DMG-PEG 2000) were dissolved in ethanol prior to formulation. The molar composition of the lipid mixture was ionizable lipid:DOPE:cholesterol:C16-PEG ceramide at 26.5:20:52:1.5 or 26.5:30:43:0.5.

RNA was diluted in a 2:1 (v/v) mixture of phosphate-buffered saline (PBS) and 10 mM citrate buffer. During formulation, the ethanol lipid phase and aqueous RNA phase were combined at a 1:3 volume ratio using a total flow rate of 12 mL/min. The resulting LNP suspension was diluted 40-fold in 1× PBS (SH30028.02, Cytiva) and concentrated by ultrafiltration using an Amicon Ultra-15 centrifugal filter unit (UFC9010).

### Characterization of LNPs

The hydrodynamic size and zeta potential of LNPs were measured after dilution to an RNA concentration of 0.001 mg/mL in 1× PBS. Dynamic light scattering (DLS) was used to determine the number-weighted mean particle size, PDI. RNA encapsulation efficiency was quantified using the Quant-iT™ RiboGreen® assay (Life Technologies).

### *In vitro* screening of LNP candidates

Raw 264.7 cells (ATCC) were seeded in clear 24-well plates at a density of 5 × 10^4^ cells per well. Lipid nanoparticles (LNPs) encapsulating EGFP mRNA were prepared using a microfluidic mixing method as previously described, with ionizable lipid candidates, DOPE, cholesterol, and PEG-lipid components. Cells were incubated with mRNA-loaded LNPs for 16 h, after which EGFP expression was quantified by measuring FITC fluorescence using flow cytometry (MA900, SONY).

### *In vivo* screening of LNP candidates

All animal experiments were conducted in accordance with protocols approved by the Institutional Animal Care and Use Committee of Seoul National University. For *in vivo* evaluation of LNP candidates, LNPs encapsulating fLuc mRNA were prepared as described above and administered intramuscularly to mice at a dose of 0.05 mg/kg (fLuc mRNA). Six hours after injection, D-luciferin was administered intraperitoneally and allowed to distribute for 15 min prior to imaging. Luciferase expression was assessed by whole-body bioluminescence imaging and *ex vivo* imaging of harvested organs using an IVIS system (PerkinElmer).

### *In vivo* cytokine and chemokine analysis

Seven-week-old BALB/c mice (Orient Bio) were intramuscularly injected with LNPs encapsulating SARS-CoV-2 spike protein mRNA at a dose of 0.5 mg/kg. Blood samples were collected at indicated time points via cheek bleeding, and serum was isolated by centrifugation. MCP-1 levels were quantified using a mouse MCP-1 ELISA kit to assess early immune responses. In addition, IL-6 concentrations were measured at the same time points using a mouse IL-6 ELISA kit according to the manufacturer’s instructions.

### Lipid Fusion FRET Assay

A FRET assay was performed to evaluate lipid fusion between LNPs and liposomes mimicking endosomal or plasma membrane compositions [25]. DOPE-conjugated FRET probes, NBD-PE and N-Rh-PE, were incorporated into liposomes, where N-Rh-PE served as a quencher of NBD fluorescence. Endosome-mimicking liposomes were prepared using BMP, DOPC, DOPE, NBD-PE, N-Rh-PE, and PI at a molar ratio of 10:50:18:1:1:10, whereas plasma membrane–mimicking liposomes were composed of DOPC, DOPE, NBD-PE, N-Rh-PE, sphingomyelin, and cholesterol at a molar ratio of 20:18:1:1:20:30.

In a 96-well plate, 100 µL of PBS (pH 7.4 or 5.5) was mixed with 1 µL of liposomes (1 mM) and LNPs at a liposome-to-LNP ratio of 1:10. Untreated wells (F_min_) and wells treated with Triton X-100 (F_max_) were used as negative and positive controls, respectively. Samples were incubated at 37 °C, and fluorescence was monitored from 5 to 90 min using a plate reader (excitation: 465 nm; emission: 520 nm). Lipid fusion efficiency was calculated as:

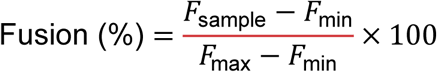

### Vaccination Schedule and Sample Collection

BALB/c mice were immunized intramuscularly with LNP formulations encapsulating SARS-CoV-2 spike mRNA (0.25 mg/kg) following a prime–boost regimen (day 0 and day 21). For cellular immune profiling, mice (n = 4 per group) were sacrificed at day 35 (two weeks post-boost), and splenocytes and draining lymph node cells were harvested for intracellular cytokine staining and flow cytometric analysis of Tfh and germinal center B cell populations. For antibody analysis, blood samples were collected at day 42 (three weeks post-boost) from a separate cohort of mice (n = 4 per group), and serum was isolated for IgG quantification and neutralization assays.

### Flow Cytometry and Intracellular Cytokine Staining (ICS)

Single-cell suspensions were prepared from the spleens and draining lymph nodes of vaccinated mice by mechanical dissociation through a 70-μm cell strainer. For intracellular cytokine analysis, cells were stimulated *ex vivo* with a SARS-CoV-2 Wuhan wild-type spike peptide pool (Miltenyi Biotec) in the presence of the protein transport inhibitors GolgiStop and GolgiPlug (BD Biosciences) for 5 h at 37°C in 5% CO_2_. Following stimulation, cells were first stained for surface markers and then fixed and permeabilized using the Cytofix/Cytoperm kit (BD Biosciences). Cells were subsequently washed with Perm/Wash buffer (BD Biosciences) and stained intracellularly for cytokines. The following fluorochrome-conjugated anti-mouse antibodies were used for flow cytometry: CD8 (clone 53-6.7), CD4 (clone RM4-5), CD44 (clone IM7), IFN-γ (clone XMG1.2), TNF-α (clone MP6-XT22), IL-2 (clone JES6-5H4), CXCR5 (clone L138D7), PD-1 (clone 29F.1A12), CD19 (clone 6D5), CD3 (clone 17A2), CD95 (clone DX2), GL7 (clone GL7), and CD62L (clone MEL-14). Dead cells were excluded using the Zombie Aqua Fixable Viability Kit (BioLegend). Data were acquired on a CytoFLEX flow cytometer (Beckman Coulter) and analyzed using FlowJo software (Tree Star). Cytokine-producing T cells were quantified as the percentage of total CD4⁺ or CD8⁺ T cells.

### Enzyme-Linked Immunosorbent Assay (ELISA)

96-well high-binding EIA/RIA plates (Costar) were coated overnight at 4°C with recombinant SARS-CoV-2 spike protein at a concentration of 2 μg/mL in phosphate-buffered saline (PBS, Welgene). Plates were then washed three times with 200 μL per well of PBS containing 0.05% Tween 20 (Sigma-Aldrich) (PBS-T) and blocked with blocking buffer consisting of 1% blotting-grade blocker (Bio-Rad) in PBS-T for 1 h at 37°C. Mouse serum samples were serially diluted 4-fold in blocking buffer, added to the plates, and incubated for 2 h at room temperature. Following three washes with PBS-T, horseradish peroxidase (HRP)-conjugated goat anti-mouse IgG (SouthernBiotech) diluted in blocking buffer was added and incubated for 1.5 h at room temperature. Plates were washed three additional times with PBS-T, and bound antibodies were detected using TMB substrate solution (OptEIA reagent set, BD Biosciences). The reaction was terminated by adding 0.5 M hydrochloric acid, and absorbance was measured at 450 nm using a microplate reader (Victor 3, PerkinElmer).

### Plaque Reduction Neutralization Test (PRNT)

Neutralizing antibody responses were evaluated by plaque reduction neutralization test (PRNT). PRNT₅₀ titers were defined as the reciprocal of highest serum dilution resulting in >50% reduction in plaque numbers compared with virus-only controls. Serum samples collected 5 weeks after the first immunization were heat-inactivated at 56 °C for 30 min prior to use.

Sera were serially diluted (1:10–1:5120) in serum-free medium and incubated with SARS-CoV-2 (50 PFU for mouse serum) at 37 °C for 1 h. The virus–serum mixtures were then added to Vero E6 cells seeded in 12-well plates (2 × 10⁵ cells per well) and incubated for 1 h at 37 °C under 5% CO₂. Following virus adsorption, cells were overlaid with 1.2% agar-containing medium and incubated for 2 days. Plaques were visualized by crystal violet staining and counted to determine PRNT₅₀ titers.

### Microneutralization assay

Vero cells were seeded at 1.5 × 10^4^ cells per well in Dulbecco’s Modified Eagle’s Medium (DMEM, Welgene) supplemented with 1% fetal bovine serum (FBS, Gibco) in 96-well plates (SPL) 24 h prior to infection. Mouse serum samples were serially 2-fold diluted in triplicate for eight dilution steps, starting from the initial dilution, and mixed with an equal volume of SARS-CoV-2 Wuhan wild-type containing 100 tissue culture infective dose 50% (TCID50). Serum–virus mixtures were pre-incubated for 30 min at room temperature and then transferred onto the Vero cell monolayers. After 96 h of incubation at 37 °C in a 5% CO_2_ environment, cells were fixed by adding paraformaldehyde (Curebio) to a final concentration of 4% and incubated for 10 min at room temperature. Wells were washed twice with 200 μL PBS per well, and the fixed cell monolayers were stained with 0.1% crystal violet solution for 10 min. After staining, wells were washed three times with 200 μL distilled water per well. Virus-induced cytopathic effects were evaluated by bright-field visual inspection of the stained cell monolayers. Serum neutralizing antibody titers were determined as endpoint MN50 values, defined as the reciprocal serum dilution conferring ≥50% neutralization, with titers between two dilution points interpolated using the geometric mean.

### IFN-γ enzyme-linked immunospot assay (ELISpot)

The assay was performed using a mouse IFN-γ ELISpot kit (XEL485, R&D Systems) according to the manufacturer’s instructions. Splenocytes (2.5 × 10⁵ cells per well) were plated in 96-well PVDF plates pre-coated with anti–mouse IFN-γ capture antibody and stimulated with SARS-CoV-2 spike glycoprotein (100 ng/well, GeneScript). Wells containing medium alone served as negative controls, and a cell stimulation cocktail (eBioscience) was used as a positive control. Plates were incubated at 37 °C for 18–24 h.

After incubation, plates were washed and incubated with a biotinylated anti–mouse IFN-γ detection antibody, followed by streptavidin–alkaline phosphatase. Spots were developed using BCIP/NBT substrate and quantified using an ImmunoSpot reader.

### SARS-CoV-2 Challenge Study

Challenge experiments were performed under biosafety level 3 (BSL-3) conditions in accordance with institutional guidelines and protocols approved by the Institutional Animal Care and Use Committee of the Korea Centers for Disease Control and Prevention (KDCD-IACUC-22-004). K18-hACE2 transgenic mice were immunized intramuscularly with SARS-CoV-2 spike mRNA–encapsulated LNPs following a prime–boost regimen (0.25 mg/kg; 5 µg in 50 µL). Mice received a primary immunization (week 0) and a booster immunization three weeks later. Three weeks after the booster dose, animals were challenged with SARS-CoV-2 under institutionally approved biosafety conditions. Clinical signs and body weight were monitored daily after challenge. At predetermined endpoints, protection was evaluated by survival and clinical outcomes, and tissues were collected for downstream analyses as described in the institutional protocol.

### Single-cell RNA sequencing

scRNA-seq was performed to characterize early immune responses at the injection site following mRNA-LNP administration, as previously described with minor modifications [17]. BALB/c mice were intramuscularly injected with SARS-CoV-2 spike mRNA encapsulated in LNPs (5 µg in 50 µL; 0.25 mg/kg). Muscle tissues from the injection site were collected 16 h post-injection and processed to obtain single-cell suspensions.

Excised muscle tissues were mechanically dissociated and enzymatically digested, followed by filtration through a 70-µm cell strainer. Viable cells were counted and used for library preparation. Single-cell libraries were generated using a droplet-based platform (10x Genomics Chromium) according to the manufacturer’s protocol and sequenced on an Illumina platform.

Raw sequencing data were processed using the Cell Ranger pipeline for alignment and gene counting. Downstream analyses were performed using the Seurat package in R. Low-quality cells were excluded based on gene counts and mitochondrial gene content. After normalization and scaling, dimensionality reduction was performed using principal component analysis (PCA) and UMAP. Cell populations were identified using graph-based clustering and annotated according to canonical marker gene expression. Differential gene expression analysis was performed to evaluate inflammatory and chemokine responses.

### Statistical Analysis

All data are presented as mean ± standard error of the mean (SEM) unless otherwise indicated. Statistical analyses were performed using GraphPad Prism (version 9.0.2, GraphPad Software). Comparisons between two groups were conducted using an unpaired two-tailed Student’s *t*-test. For multiple group comparisons, one-way analysis of variance (ANOVA) followed by Tukey’s multiple comparisons test was used. Survival curves were analyzed using the Kaplan–Meier method and compared by the log-rank (Mantel–Cox) test when applicable. A *p* value < 0.05 was considered statistically significant. The number of biological replicates (*n*) is indicated in the corresponding figure legends.

### Use of Artificial Intelligence

Manuscript drafting was assisted by Claude (Anthropic), an AI language model. All AI-generated content was critically reviewed, edited, and verified by the authors, who take full responsibility for the accuracy and integrity of the reported findings

## Acknowledgements

This work was supported by the Bio & Medical Technology Development Program of the National Research Foundation of Korea (NRF), funded by the Ministry of Science and ICT (MSIT) under the following grant numbers: RS-2023-00261343, RS-2024-00451880, RS-2024-00411768, RS-2025-00542986, RS-2024-00465298 and RS-2025-25455885, and by the Korea National Institute of Health (Grants 4800-6634-329; intramural research number: 2022-NI-001-02). Schematic figures were created with BioRender.com

## Data Availability Statement

The data that support the findings of this study are available from the corresponding author upon reasonable request.

## Notes

### Competing Interest Statement

E.H.J., J.H.K., M.S.K., H.E.H., O.H.K., S.K. are employees of Surginex Co., Ltd. The remaining authors declare no competing interests.

