## Supplementary Data for "Coordinated Tuning of Ionizable Lipids and Formulation Redirects mRNA Vaccines Toward Lymphoid-Specific CD4^+^ T Cell Immunity"

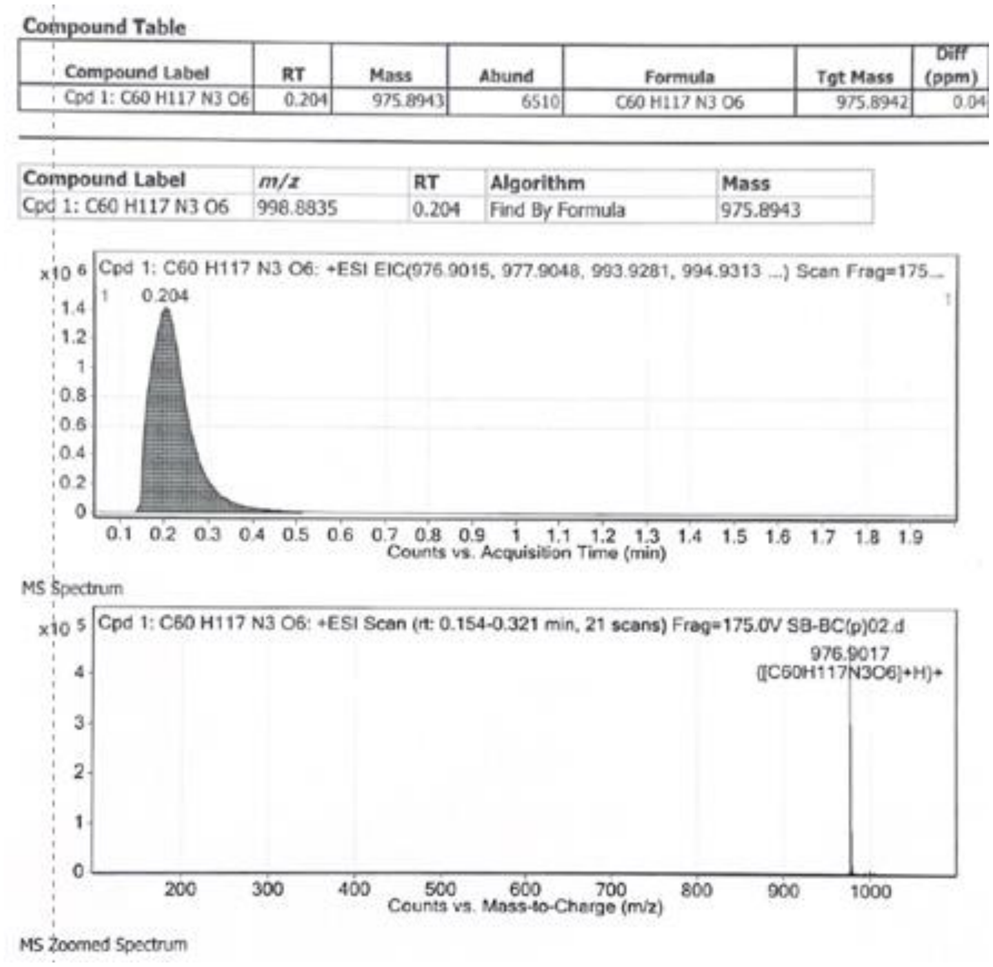


**Figure S1. LC–MS of N4Z.** A single peak at RT = 0.204 min and a major ion at *m/z* 976.90 [M+H]+[M+H]^+[M+H]+ confirm the identity and purity of N4Z.

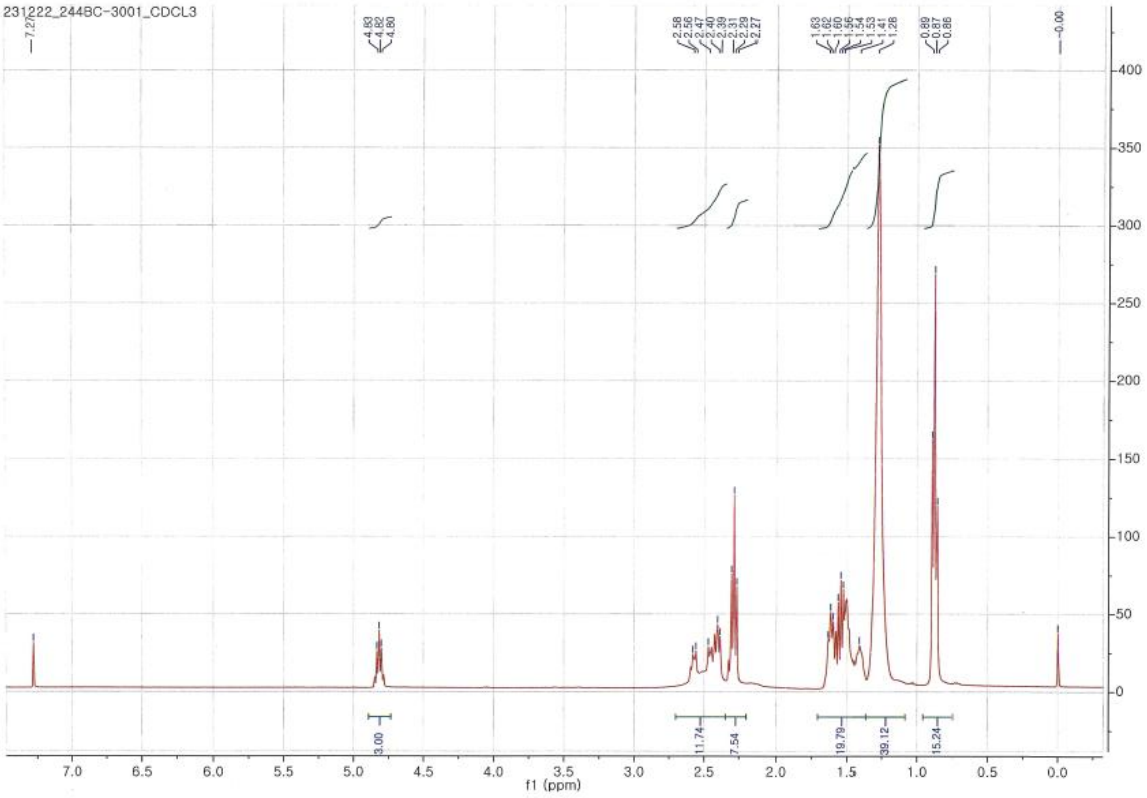


**Figure S2. ¹H NMR spectrum of N4Z.** The ¹H NMR spectrum ($CDCl_3$) confirms the chemical structure of N4Z, showing characteristic proton signals consistent with the expected molecular structure.

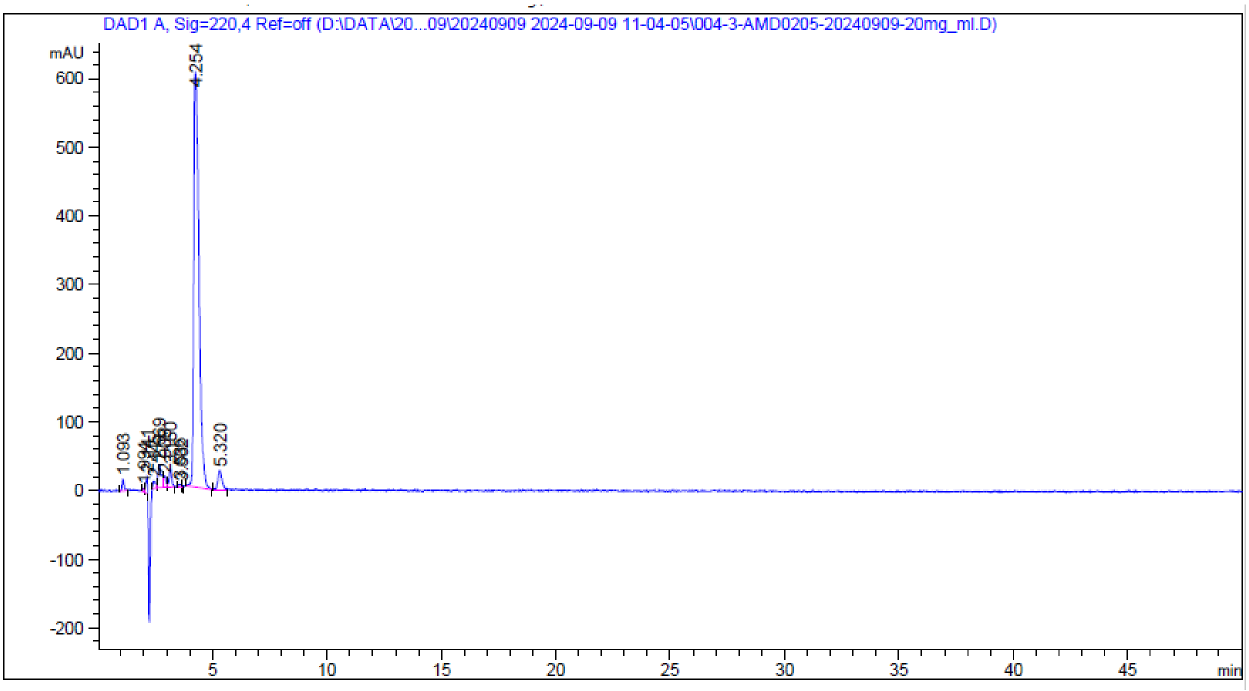


**Figure S3. HPLC chromatogram of 244-BC.**

A major peak at RT = 4.254 min (86.7%) confirms the target compound, with a minor impurity peak at RT = 5.320 min (3.21%).

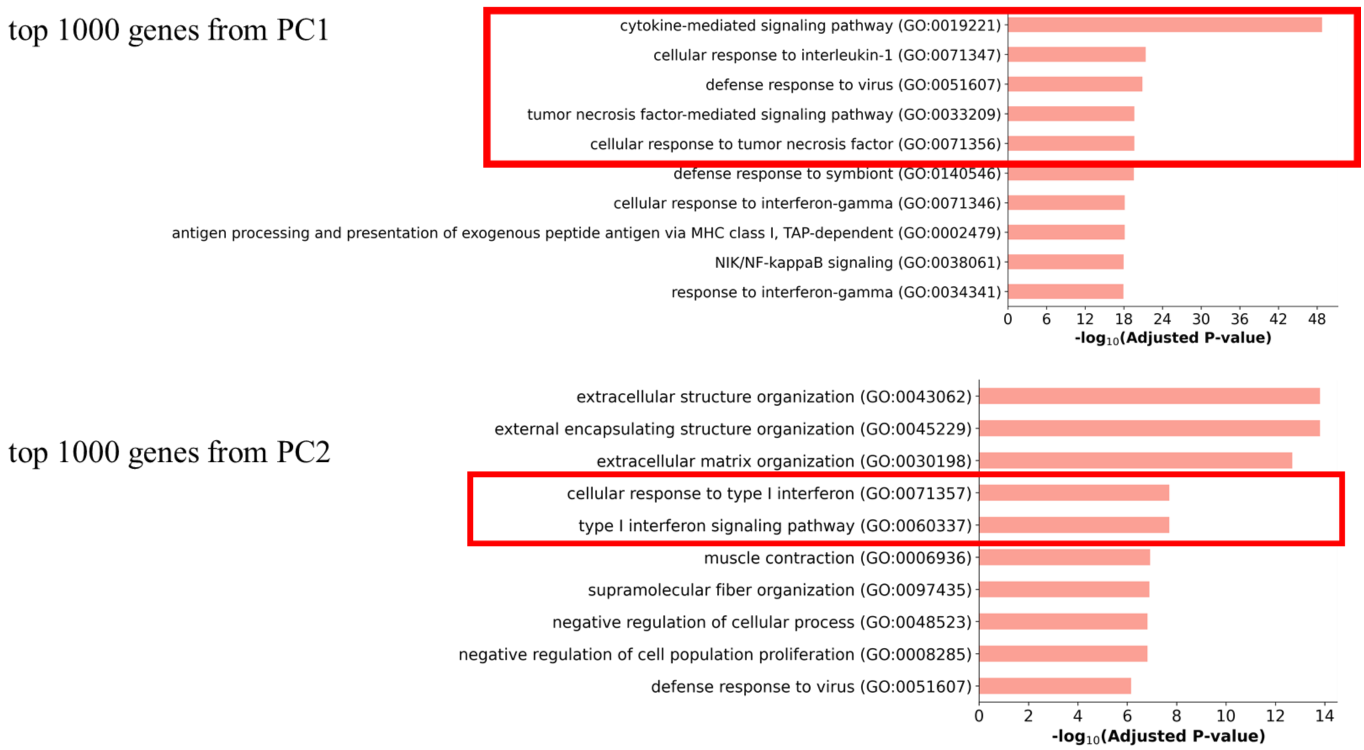


**Figure S4. Pathway enrichment of principal component–associated gene programs.** Gene ontology analysis of the top genes contributing to each principal component shows that PC1 is associated with inflammatory and cytokine signaling pathways, whereas PC2 is enriched for type I interferon and antiviral response pathways. Significance is shown as $-\log_{10}$ (adjusted *P* value).

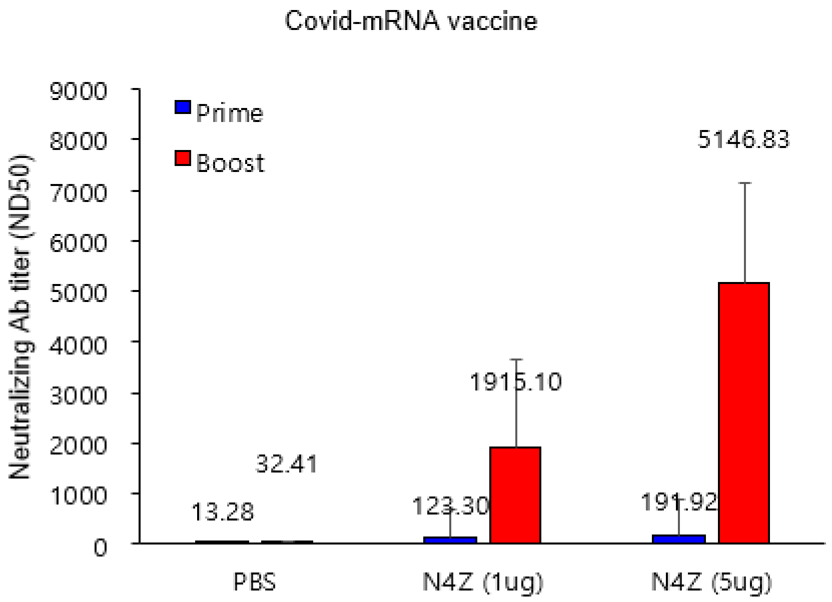


**Figure S5. Neutralizing antibody titers after N4Z vaccination.** Booster immunization significantly increased neutralizing antibody responses in a dose-dependent manner.

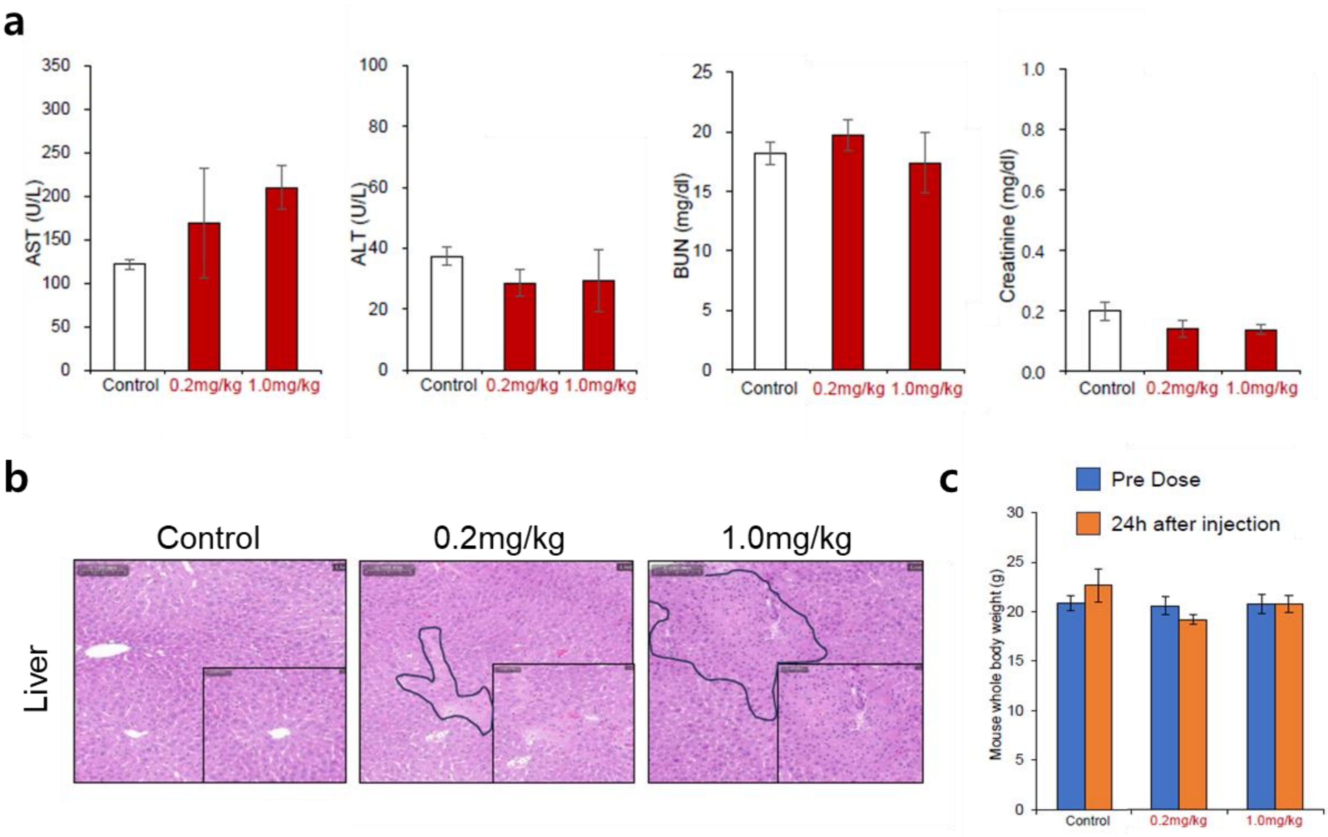


**Figure S6. Safety evaluation of N4Z lipid nanoparticles in BALB/c mice.** BALB/c mice were intramuscularly administered N4Z LNPs at 0.2 or 1.0 mg/kg, and safety parameters were assessed 24 h post-injection. (a) Serum liver function markers (AST, ALT) and kidney function markers (BUN, creatinine). (b) Representative hematoxylin and eosin (H&E)-stained liver sections from control and N4Z-treated mice. (c) Mouse whole body weight measured before (pre-dose) and 24 h after injection.

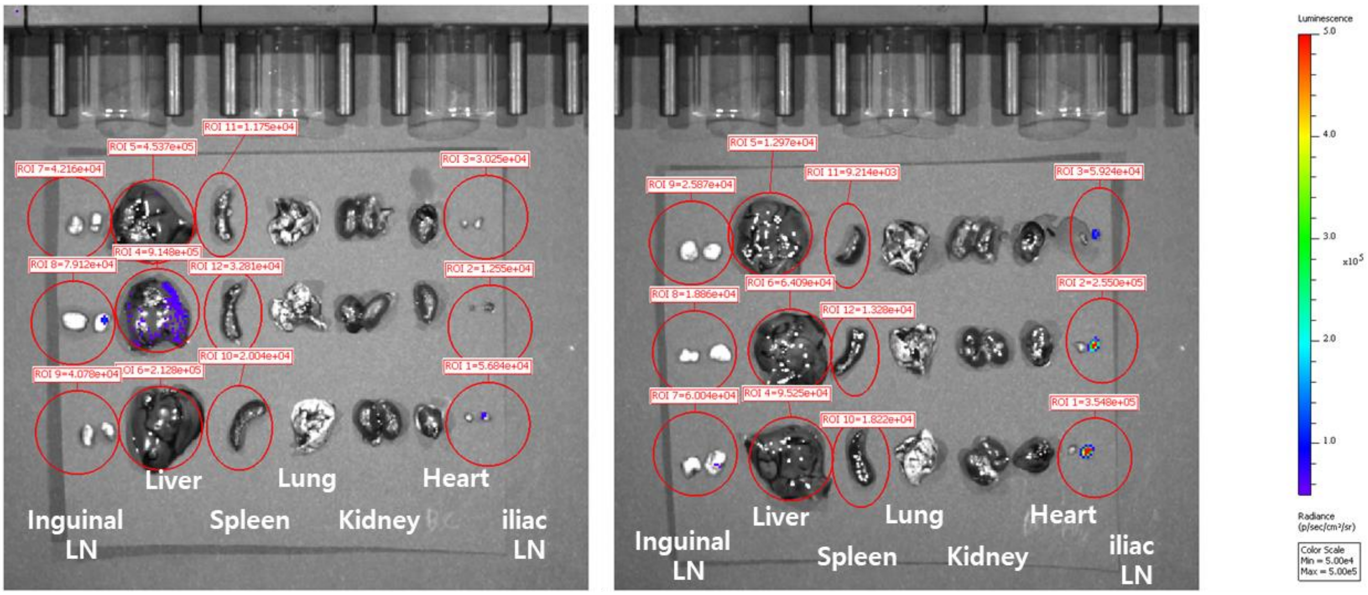


N4Z N4Z-opt

**Figure S7. Ex vivo organ biodistribution following intramuscular administration.** Representative ex vivo bioluminescence images of harvested organs (inguinal lymph node, liver, spleen, lung, kidney, heart, and iliac lymph node). Luminescence signals indicate predominant expression at the injection-draining lymph node and liver, with minimal signal in other organs.

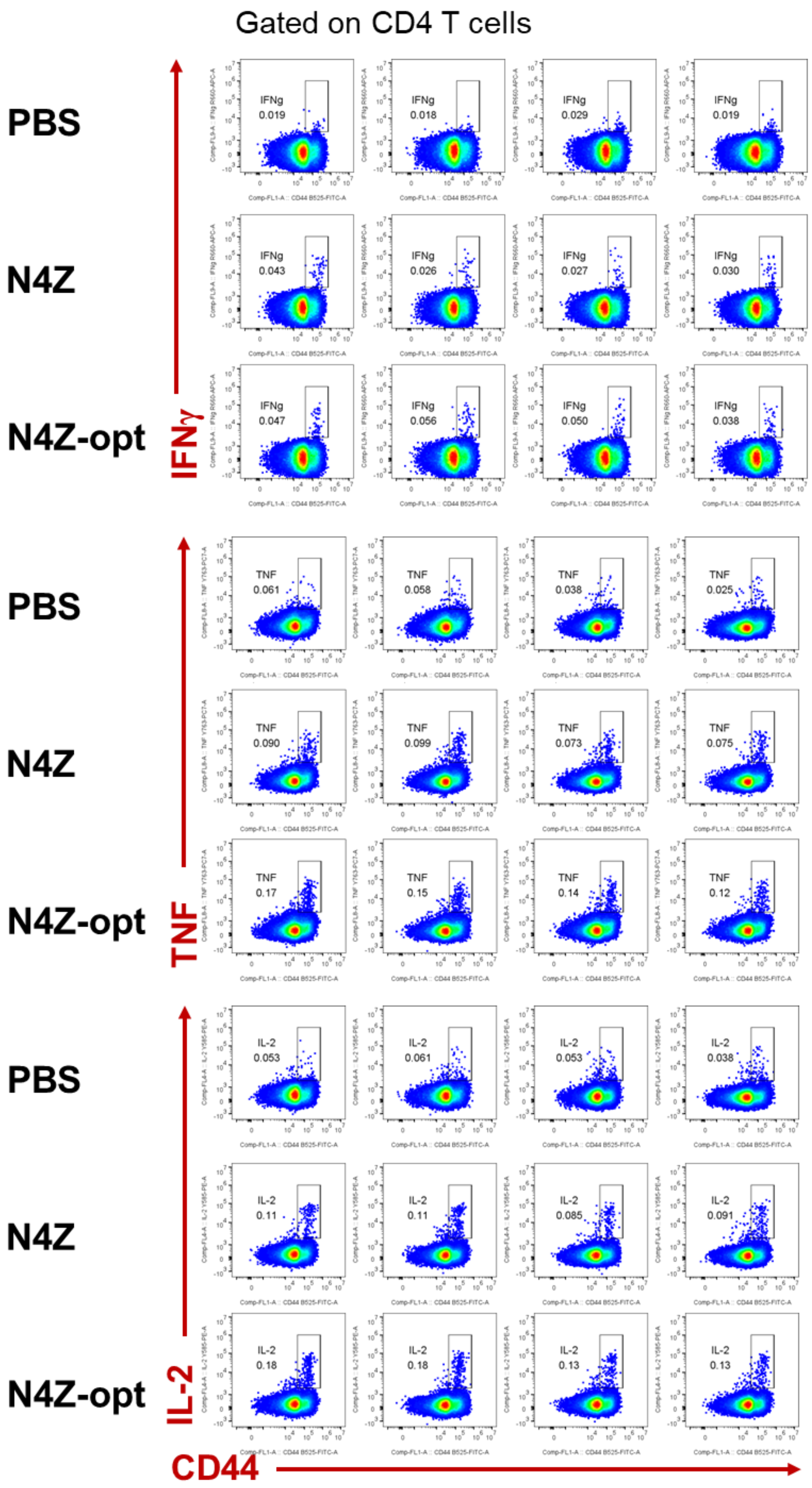


**Figure S8. Representative flow cytometry plots of splenic cytokine-producing CD4⁺ T cells.** Dot plots show IFN-γ⁺, TNF-α⁺, and IL-2⁺ CD4⁺ T cells from spleen gated on CD44⁺ cells after vaccination.

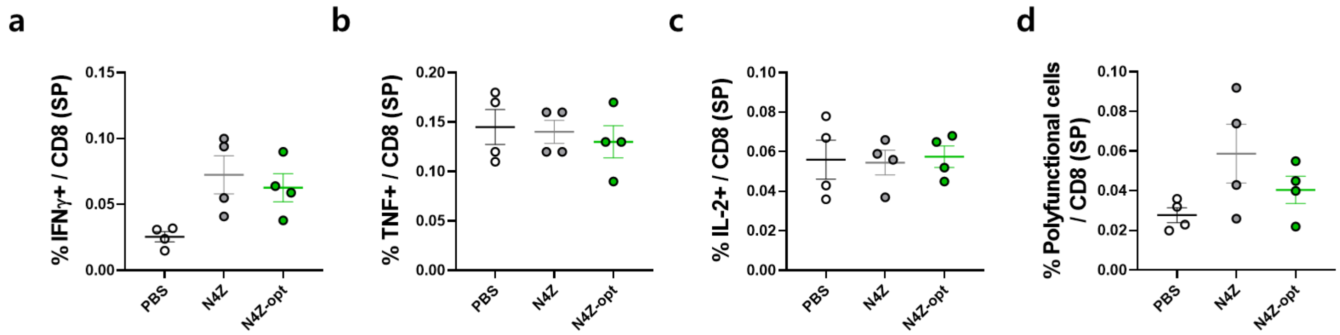


**Figure S9. CD8⁺ T cell responses (N4Z vs N4Z-opt).** N4Z and N4Z-opt induced comparable functional CD8⁺ T cell responses (mean ± SEM).

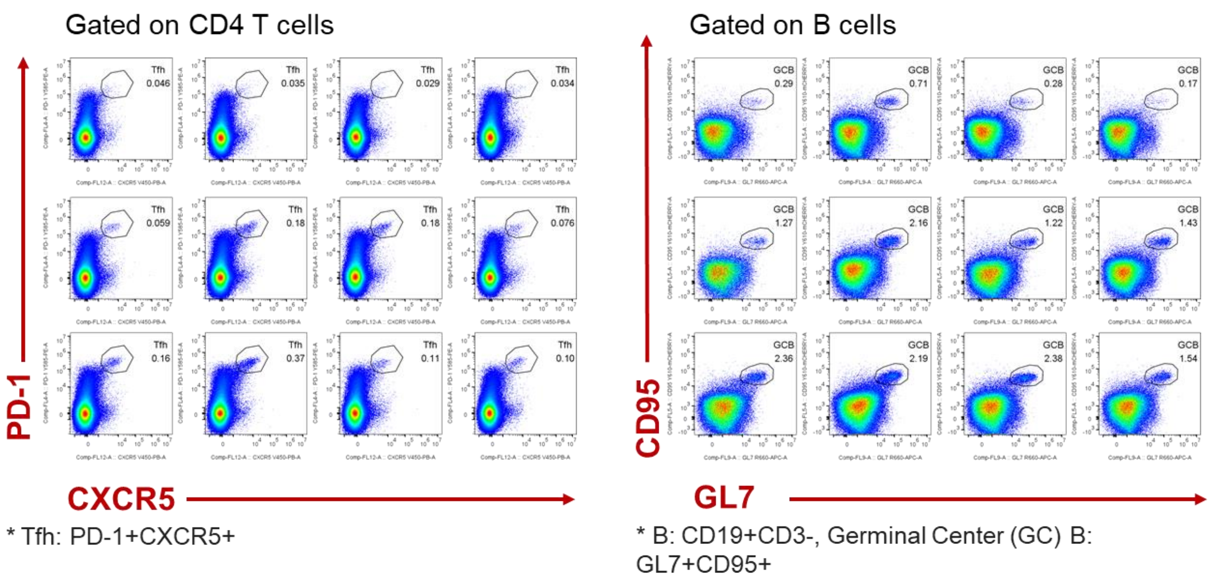


**Figure S10. Representative flow cytometry plots of Tfh and germinal center B cells from lymph nodes.** Dot plots show Tfh (PD-1$^+$CXCR5$^+$) and GC B (CD95$^+$GL7$^+$) cell populations in draining lymph nodes after vaccination.
